# Cell-type specific inhibitory dynamics shape binocular response normalization in visual cortex

**DOI:** 10.64898/2026.09.20.753031

**Authors:** MB Perumal, H Bushnell, D Dopp, S Gharaei, E Arabzadeh, SS Nair, GJ Stuart

**Affiliations:** Department of Physiology, Biomedicine Discovery Institute, Monash University, Clayton, Australia; Division of Neuroscience, John Curtin School of Medical Research, Australian National University, Canberra, Australia; Department of Electrical Engineering and Computer Science, University of Missouri, Columbia, United States of America

**Keywords:** inhibition-stabilized network, normalization, binocular summation, somatostatin interneurons, parvalbumin interneurons, visual cortex, inhibition

## Abstract

Response normalization during sensory processing is a canonical cortical computation thought to emerge from inhibition-stabilized networks (ISNs). A key prediction from ISNs is that inhibition tracks and scales with excitation, leading to sublinear response summation, but which interneuron classes mediate this operation is unclear. Here, we investigate temporal dynamics and response summation in parvalbumin (PV) and somatostatin (SST) interneurons in mouse binocular visual cortex (bV1). We find that both PV and SST interneurons are binocular but exhibit distinct temporal dynamics and summation profiles. Binocular responses in PV interneurons tracked pyramidal neuron responses, exhibiting sublinear summation consistent with ISN models. In contrast, binocular responses SST interneurons exhibited rapid temporal dynamics and linear or supralinear summation, deviating from ISN predictions. While both PV and SST neurons received excitatory drive via interhemispheric callosal input, they exhibited distinct cellular properties, synaptic dynamics and input-output transformations. Biophysically constrained simulations indicated that cellular and synaptic differences alone were insufficient to reproduce the observed response dynamics in vivo during binocular integration, which were better explained by class-specific differences in local inhibitory circuit motifs. Together, our results indicate that heterogeneous, cell-type-specific inhibitory dynamics shape response normalization in bV1, with PV but not SST interneurons having ISN-like properties.

## Introduction

Recurrent excitatory circuits are a ubiquitous feature of the cerebral cortex ^1,2^. Theoretical models predict that recurrent excitation can amplify weak inputs and support persistent activity^3,4^, however, such amplification risks runaway excitation unless stabilized by inhibition ^5^. The cortical computation underlying stabilisation of recurrent excitation by inhibition during sensory processing is often referred to as normalization. It allows cortical circuits to flexibly scale their responses across a wide range of stimulus strengths and input sources without saturation or runaway excitation. Normalization is observed across sensory cortices and species ^6,7^ and serves to stabilize network activity during integration of diverse inputs from multiple sources ^5,8–10^.

A core signature of normalization is sublinear response summation, in which the response to simultaneous stimuli is smaller than the sum of responses to each stimulus presented alone ^6,11^. Consistent with this, in the mouse binocular visual cortex (bV1) excitatory pyramidal neurons exhibit sublinear summation of convergent inputs from the eyes during binocular integration ^12–14^. Pharmacological perturbations and computational modelling have implicated inhibition as a central mechanism generating sublinear binocular responses in bV1 ^12,13^.

Sublinear summation also emerges in cortical models operating in inhibition-stabilized networks (ISNs) ^5,11,15,16^. In these models, inhibitory activity tracks excitation and undergoes sublinear response summation during convergent input ^11,17,18^. Cortical circuits have been proposed to operate as ISNs during sensory integration of multiple inputs ^11,15,16^. However, whether specific interneuron classes conform to, or deviate from, the inhibitory response dynamics predicted by ISN models is unclear.

In V1, parvalbumin-expressing (PV) and somatostatin-expressing (SST) interneurons represent major sources of inhibition that modulate excitatory gain and response dynamics ^9,19,20^. Here, we investigated whether PV and SST interneurons in bV1 exhibit inhibitory dynamics predicted by ISN models using binocular summation as a tractable computation. To investigate this, we recorded spiking from optically-tagged layer 2/3 PV and SST interneurons during monocular and dichoptic binocular stimulation. We find that both PV and SST interneurons are binocular, but exhibit distinct temporal dynamics and binocular response summation profiles. PV interneuron activity tracked the excitatory population response showing sublinear binocular summation, whereas SST interneurons showed rapid temporal dynamics with linear-to-supralinear summation. In vitro recordings in bV1 revealed that both PV and SST interneurons receive excitation via an interhemispheric pathway, but have different intrinsic properties, synaptic dynamics and input-output transformations. Using biophysically constrained models we show that differences in cellular and synaptic properties cannot explain the divergent temporal dynamics seen in PV and SST interneurons during binocular integration. Instead, we find that differences in PV and SST temporal dynamics are driven by cell-type specific differences in inhibitory circuit motifs. Together, our findings reveal heterogeneous, cell-type specific inhibitory properties shape cortical normalization during binocular integration.

## Results

### Monocular and binocular responses in optically tagged PV and SST units in bV1

We recorded the responses of PV and SST neurons in mouse bV1 during monocular and binocular stimulation. To stimulate each eye separately (monocular) or both eyes together (binocular), we positioned a haploscope in front of the snout (Figure 1A). Each mirror reflected high contrast sinusoidal drifting grating stimuli (12 directions, 2 Hz temporal frequency for 0.5 s) displayed on two independently controlled monitors ^12^. Recordings were performed in transgenic mouse lines expressing channelrhodopsin-2 in either PV or SST interneurons. This allowed us to identify PV and SST interneurons following optical tagging with brief blue LED illumination of the cortical surface (470 nm, 50 ms).

**Figure 1.**
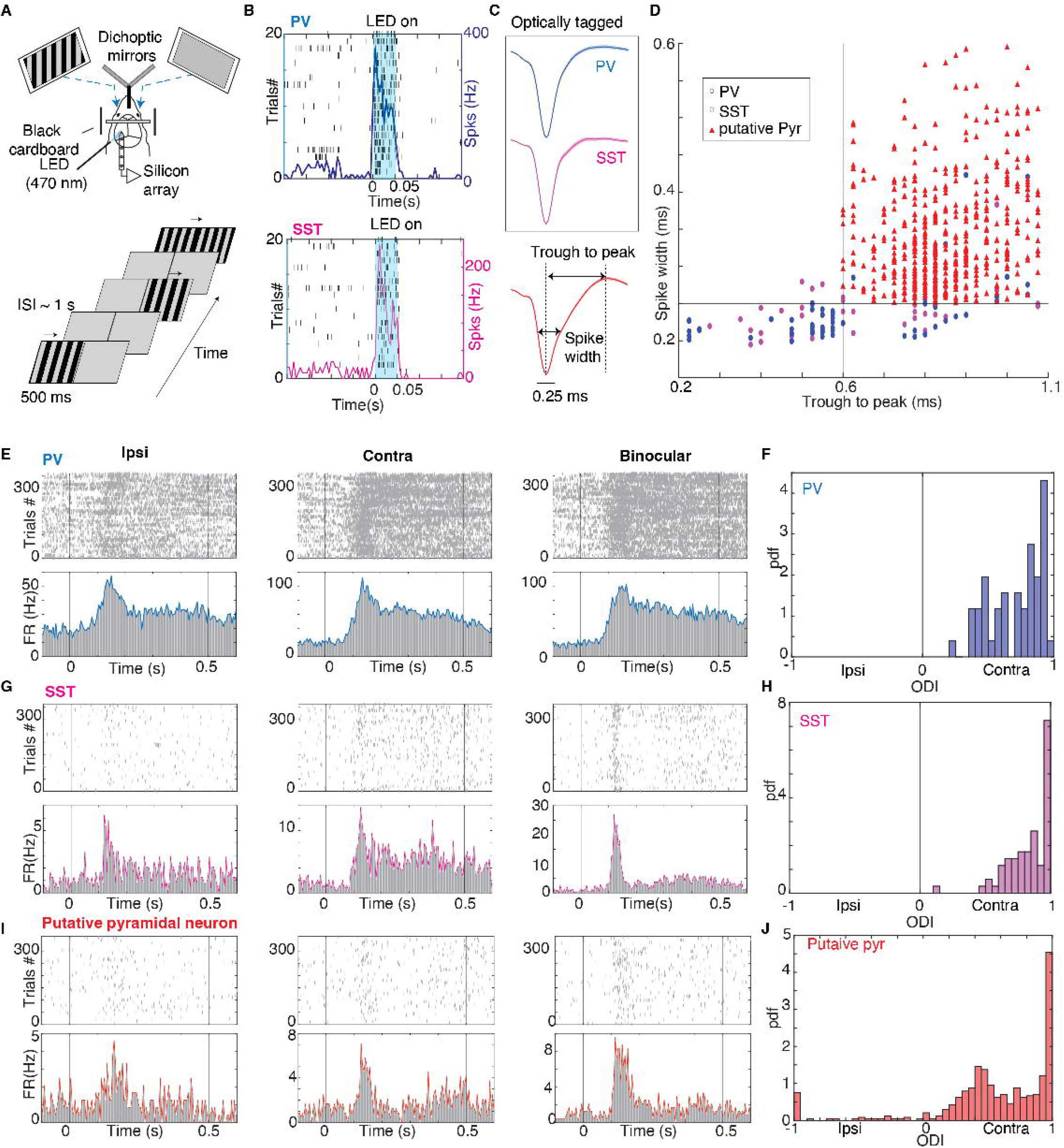
Cell-type classification and ocular dominance distributions. (A) Experimental set-up with mouse facing a haploscope for presentation of visual stimuli separately to each eye. Extracellular activity was recorded using a silicon array inserted through a cranial window over V1. A blue LED (470nm) was placed above V1 for optical tagging of neurons expressing channelrhodopsin (ChR2) in SSTCre or PVCre mice. Visual stimuli comprised of drifting gratings presented for 0.5 s with a randomized interstimulus interval between 1-1.25 s. (B) Raster plot of opto-tagged PV (top) and SST (bottom) units showing an increase in spike response during LED pulses. (C) Average waveforms of an opto-tagged PV and SST unit and a non-opto-tagged unit with broad spike features. (D) Distribution and identification of putative pyramidal units (red triangles) among non-opto-tagged units using a criterion for spike width and trough-peak distance. (E) Monocular and binocular responses in the opto-tagged PV unit. Left to right, raster plots (top) and PSTH (bottom, 5 ms bins) in response to stimulation of ipsilateral (ipsi), contralateral (contra) and both eyes (binocular). (F) Ocular dominance index distribution of opto-tagged PV units (n=51). (G) Monocular and binocular responses in an opto-tagged SST unit. Left to right, raster plots (top) and PSTH (bottom, 5 ms bins) in response to stimulation of ipsilateral (ipsi), contralateral (contra) and both eyes (binocular). (H) Distribution of ocular dominance index of opto-tagged SST units (n=69). (I) Monocular and binocular responses in the identified putative pyramidal unit. Left to right, raster plots (top) and PSTH (bottom, 5 ms bins) show response to stimulation of ipsilateral (ipsi), contralateral (contra) eye and both eyes (binocular). (J) Distribution of ocular dominance index of identified putative pyramidal units (n=475).

Optically tagged units were identified post hoc based on robust light-evoked increases in firing rate (Figure 1B) using receiver operating characteristics (ROC)-based criteria (Methods). Putative pyramidal units were identified by the absence of light-evoked responses and by broader spike waveforms, including a longer spike-width and trough-to-peak distance ^21^ (Figure 1C and D). Units with significantly increased spiking within 300 ms after visual stimulus onset were classified as visually responsive (see Methods). Overall, we identified 51 visually responsive optically-tagged PV units (5 mice, 9 recordings) and 69 visually responsive optically-tagged SST units (5 mice, 10 recordings) as well as 475 visually responsive putative pyramidal units (10 mice, 19 recordings). Both PV and SST units showed similar broadly tuned responses with an orientation selective index of less than 0.1 during binocular stimulation (mean orientation selectivity index ± SD, PV 0.08 ± 0.05; SST 0.08 ± 0.06). This observation is consistent with the idea that PV and SST interneurons in bV1 pool inputs from surrounding pyramidal neurons with different orientation tuning preferences, resulting in broadly tuned inhibition ^2,21–27^. Given their low orientation selective index, in all subsequent analysis we combined and averaged responses in PV and SST units across all stimulus orientations.

Previous two-photon imaging studies have shown that PV neurons in bV1 are binocular ^28^, but whether responses in SST neurons in bV1 are also binocular is not known. Consistent with this previous work, we found that the vast majority of PV units in bV1 (96%; 49/51) responded to both contralateral and ipsilateral eye stimulation and were classified as binocular (Figure 1E,F). In comparison, approximately half of the visually responsive SST (54%; 37/69) and pyramidal units (46%; 220/475) were binocular (Figure 1G-J). Optically tagged PV and SST units, as well as pyramidal units, rapidly increased their firing rate after stimulus onset with more robust responses to contralateral eye than ipsilateral eye stimulation, consistent with prior reports of contralateral bias in mouse bV1 ^29–32^. Consistent with this, ocular dominance was greater than zero in all three cell types (Figure 1F,H&J).

To evaluate PV and SST response dynamics to convergent input from both eyes, we focused subsequent analysis on binocular units. Among binocular units, PV and SST populations showed similar ocular dominance (Figure 1F-H; PV ODI ± SD: 0.70±0.19 vs SST ODI ± SD: 0.77±0.14; Kolmogorov-Smirnov (KS) test, p=0.4). The ODI of binocular pyramidal population was significantly lower than both PV and SST population (Figure 1J ODI ± SD: 0.5±0.2; KS test, p < 0.0001), indicating the pyramidal neuron population was significantly more binocular that the PV or SST populations.

### PV and SST exhibit distinct temporal evolution of spiking responses

ISN models predict that inhibitory activity should track excitatory activity during convergent input ^11,18,33^. To evaluate the temporal dynamics of spiking responses in pyramidal, PV and SST neurons in bV1, we generated post-stimulus time histograms (PSTH; 5 ms bins) for each neuronal cell type. PSTHs were baseline-subtracted and normalized to the mean firing rate of each unit across the recording session, excluding optical tagging trials ^21,34^. This yielded a unitless measure of the proportional change in spiking, allowing comparison of the temporal dynamics across units independent of differences in overall firing rate.

In V1, incoming visual inputs via feedforward and local recurrent circuits are proposed to drive early responses (within the first 200 ms), whereas feedback inputs are thought to modulate later responses (after approximately 250 ms) ^35,36^. We therefore defined an early response window of 200 ms after the stimulus onset to evaluate response dynamics driven primarily by feedforward and local recurrent visual input.

Average responses in pyramidal, PV and SST populations rose steeply, after stimulus onset, reaching their peak within this early response window (Figure 2A). To quantify early response dynamics, we estimated the peak latency and rise time (10% to peak) during ipsilateral, contralateral and binocular stimulation. During ipsilateral eye stimulation, peak latency and rise time were comparable across PV, SST and pyramidal neuron population (Figure 2B,C; left), but these dynamics diverged during contralateral and binocular eye stimulation (Figure 2B,C; middle and right). During binocular eye stimulation, the peak of PV responses was slightly delayed relative to the pyramidal neuron population, with PV and pyramidal units having similar rise times. In contrast, during binocular eye stimulation SST responses peaked significantly earlier than both PV and pyramidal responses and exhibited faster rise times (Figure 2B,C; middle and right). These findings indicate that during weaker ipsilateral eye responses, early response timing of PV and SST activity were similar to pyramidal population activity, whereas PV and SST response dynamics diverged during stronger responses evoked by contralateral and binocular eye stimulation. Under these conditions of more powerful activation, PV responses closely tracked pyramidal population activity, whereas SST responses diverged, rising faster and peaking earlier.

**Figure 2.**
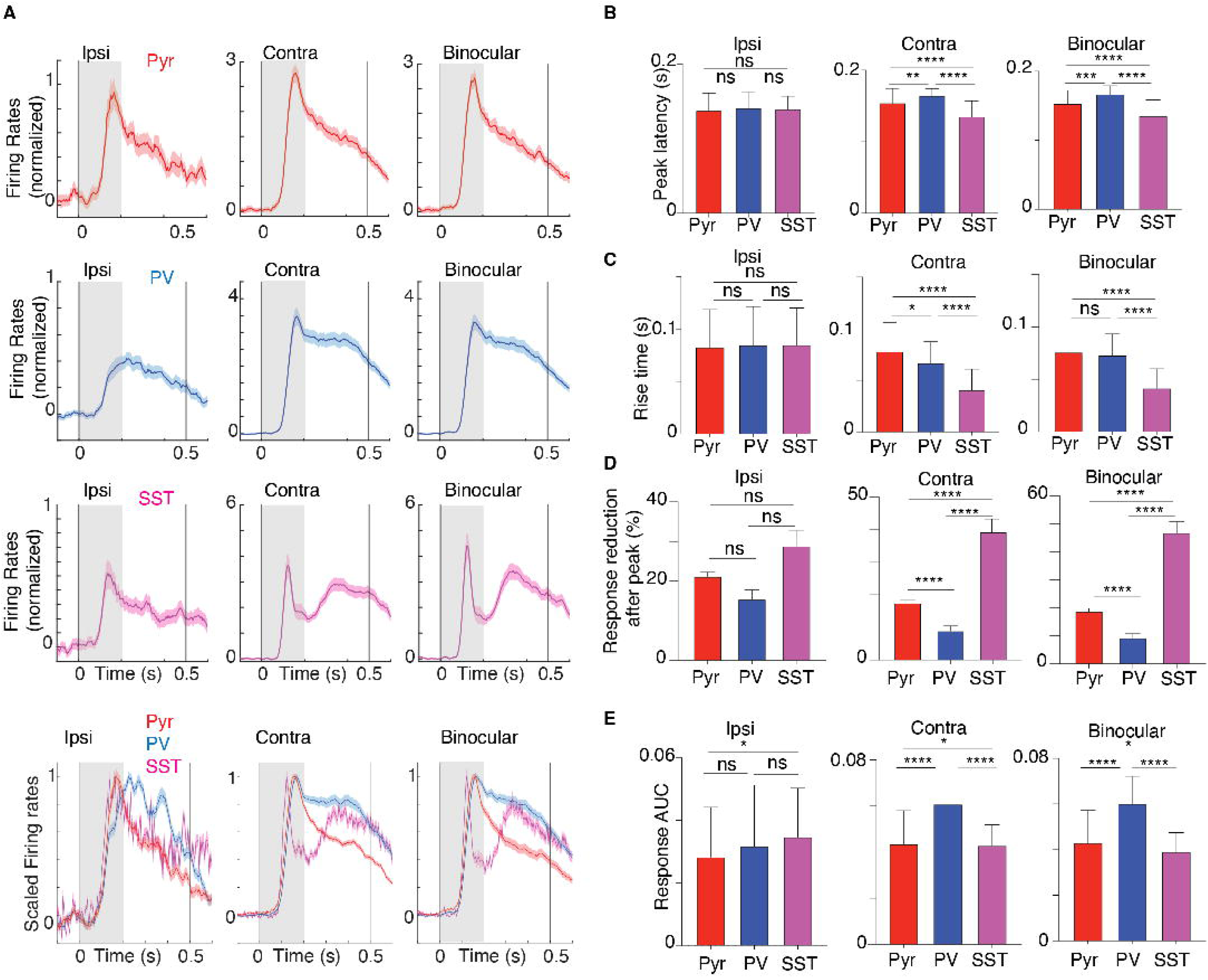
Temporal features of visual evoked responses in putative pyramidal, PV and SST units. (A) From left to right, population average ± SEM of normalized spike responses to ipsilateral (ipsi; left), contralateral (contra; middle) and binocular (right) stimulation for binocular units. From top, putative pyramidal (red), PV (blue), and SST units (magenta). Bottom, overlaid responses of putative pyramidal (red), PV (blue) and SST (magenta) scaled to peak. Early response window highlighted in grey. (B) Left to right, comparison of peak latencies (mean ± sem) of ipsi (left), contra (middle) and binocular (right) responses for pyramidal (red) PV (blue) and SST (magenta). (C) Comparison of rise times (10% to peak, mean± sd) of ipsi (left), contra (middle) and binocular (right). (D) Percentage change in spike response, calculated from the peak response to the mean post-peak response within a 50 ms window, during contralateral (left) and binocular stimulation (right). Data are shown as mean ± SEM. (E) Comparison of peak-normalized area under the curve (AUC, mean ± sd) in the early response window (0 to 200 ms from stimulus onset) for ipsi (left), contra (middle) and binocular (right) responses. Comparisons using KS test and statistical significance indicated using asterisk (*).

This fast-activating SST response is notable because SST neurons are thought to receive significantly less feedforward input compared to PV and pyramidal neurons^37,38^. Because binocular PV and SST units exhibited similar ocular dominance, the earlier peak latency and faster rise time of SST responses are unlikely to reflect a difference in eye preference. This rapid SST response under our stimulus conditions contrasts with delayed SST spiking during monocular small drifting-bar stimuli ^39^ and sparse-noise stimulation ^40^, but is consistent with SST responses observed by others during full-field, high-contrast gratings and flashes^10,38,41^.

Following the peak response, PV and pyramidal responses declined gradually, whereas SST responses decayed rapidly before recovering during later responses (>250 ms) during contralateral and binocular eye stimulation (Figure 2A). We quantified the post-peak reduction in firing as the difference between the peak firing rate and mean firing rate in a time-window 50 ms after the peak. During ipsilateral eye stimulation, the post-peak reduction in spiking was comparable across the three populations (Figure 2D, left), but diverged during contralateral and binocular eye stimulation. During binocular stimulation PV and pyramidal responses declined moderately by 9% and 19%, respectively, whereas the SST population exhibited a steep post-peak reduction of 47% (Figure 2D, middle and right). Following normalisation of responses to their peak, the area under early responses (0 to 200 ms after stimulus onset) was comparable across the three populations during ipsilateral stimulation, but was greater in PV units than in both pyramidal and SST population during contralateral and binocular stimulation (Figure 2E), indicating PV responses were more sustained than both pyramidal and SST responses.

Together, these results indicate that PV and SST populations respond with distinct temporal dynamics to binocular visual stimulation. The peak latency and rise time of PV responses closely followed pyramidal responses and was sustained, whereas SST responses rose more rapidly, peaked earlier and exhibited a steep post-peak reduction deviating from ISN dynamics. The divergent temporal profiles of PV and SST responses suggest that different cell-type specific mechanisms are engaged during strong visual drive of convergent binocular inputs.

### PV and SST neurons exhibit contrasting binocular response summation

Both excitatory and inhibitory populations in ISN models exhibit sublinear response summation during convergent input. To quantify binocular summation, for each unit we computed the difference between the binocular (B) response and the monocular sum of contralateral (C) and ipsilateral (I) responses (B-(C+I)). In pyramidal units, the population-averaged difference between the binocular response and the monocular sum was below zero throughout the visually evoked response, indicating sublinear summation (Figure 3A,B). Consistent with this, plotting the peak binocular response against the corresponding monocular sum for each unit individually indicated that the vast majority of pyramidal units (93%) fell below the unity line (Figure 3C). We then estimated a summation index for each unit as the peak binocular response minus the monocular sum, divided by the sum of these quantities ((B-(C+I))/(B+C+I)). For the pyramidal neuron population, the average summation index was significantly below zero, indicating sublinear binocular response summation (Figure 3D). These findings are consistent with previous studies showing normalization of binocular responses in pyramidal neurons through sublinear summation ^13,14^.

**Figure 3.**
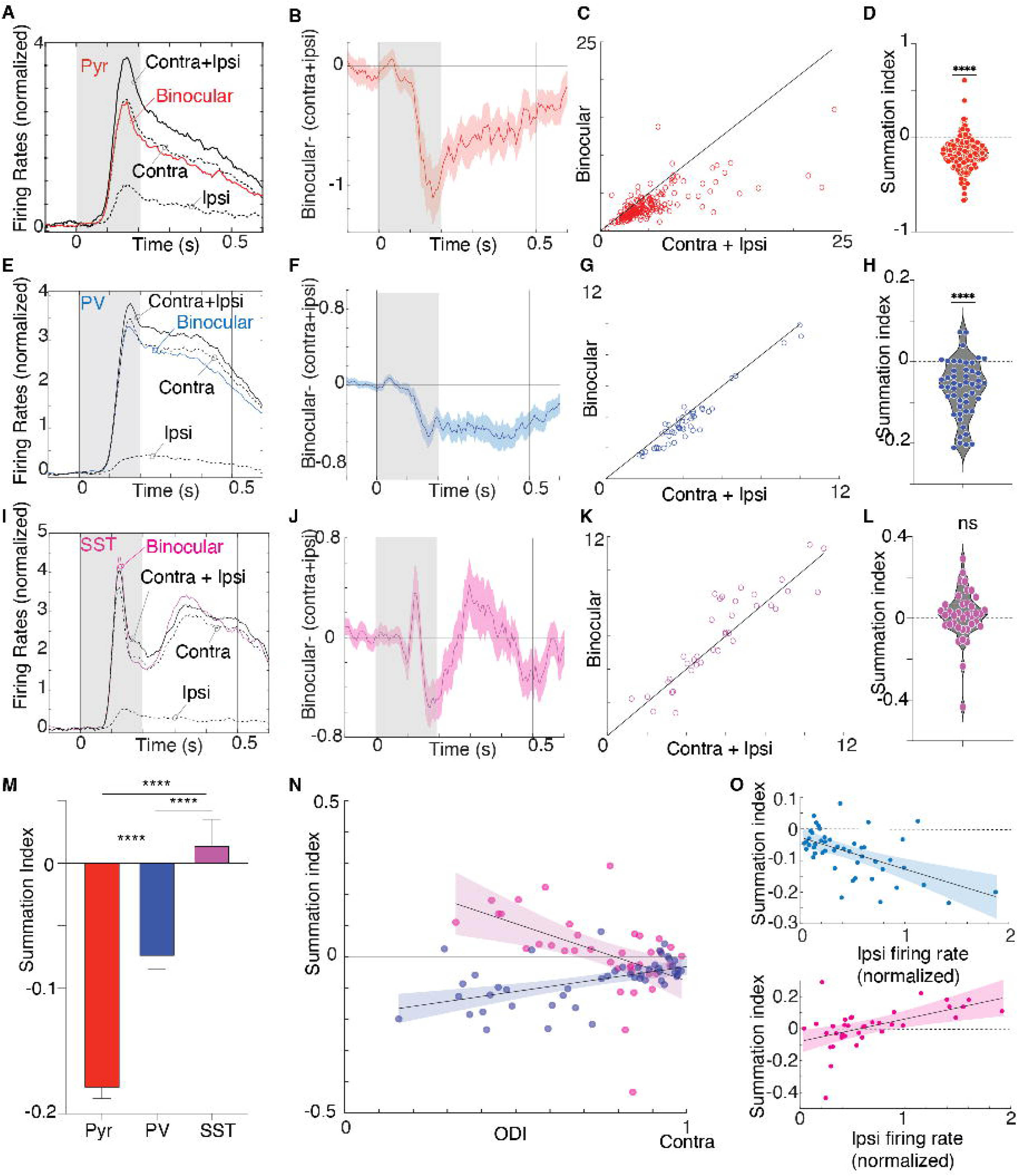
Binocular response summation in PV and SST units and relationship to ocular dominance. (A) Overlaid mean response in putative pyramidal units to ipsilateral (ipsi), contralateral (contra), binocular, and the linear sum of ipsi and contra responses. (B) Difference (mean±sem) between the binocular and linear sum of monocular responses in putative pyramidal units. (C) Peak binocular response versus linear sum of monocular responses in individual putative pyramidal units. (D) Summation index in putative pyramidal units (one sample t-test). (E-H) Same as A-D, but for PV units. (I-L) Same as in A-D, but for SST units. (M) Comparison of the average summation indices for PV, SST and putative pyramidal (Pyr) units (KS test). (N) Correlation between the summation indices for PV (blue) and SST (magenta) units and the ODI. (O) Correlation between the summation index and the ipsilateral response peak; PV (top, Pearson r =0.306, ****) and SST (bottom, Pearson r =0.3, ***).

Similar to pyramidal units, PV units showed sublinear binocular response summation throughout the duration of visual stimulation (Figure 3E,F). As with the pyramidal population, plotting the peak binocular response against the monocular sum indicated that the vast majority of PV units (86%) fell below the unit line (Figure 3G) and that the average summation index was significantly below zero, indicating sublinear summation (Figure 3H). In contrast, SST units showed near-linear or supralinear integration during the early phase of the visual response. The peak binocular response in SST units was on average greater than the monocular sum (Figure 3I). Consistent with this, the population average of the binocular response minus the linear sum showed a time-dependent change with supralinear summation early (<200 ms after stimulus onset) and late (>250 ms), separated by a period of sublinear summation (Figure 3J). Plotting the peak binocular response against the corresponding monocular sum for each SST unit individually indicated that the majority of SST units (62%) lay above the unity line, indicating supralinear summation (Figure 3K). Across all SST units the summation index of the peak SST response was, however, not statistically different from zero, indicating on average linear summation (Figure 3L). Overall, the data indicate that binocular summation in pyramidal and PV neurons was significantly more sublinear than in SST neurons (Figure 3M).

We next investigated how binocular summation relates to the ODI by plotting the ODI against the summation index for each unit (Figure 3N). In both PV and SST units, lower ODI values, reflecting more binocular responses, were associated with greater changes in summation, but in opposite directions. In PV units, the summation index positively correlated with the ODI. In contrast, SST units showed a significant negative correlation between ODI and summation index (Figure 3N). These results indicate that ODI correlates differently with binocular summation in PV and SST populations.

Finally, we investigated how the ipsilateral response magnitude impacts on binocular summation. Increases in ipsilateral response magnitude correlated with increased sublinear summation in PV and supra-linear summation in SST units (Figure 3O). This indicates that binocular response summation in both PV and SST interneurons is strongly enhanced by input from the ipsilateral eye.

Together, these findings indicate that in addition to divergent temporal dynamics, during binocular stimulation PV and SST neurons also show different and divergent response summation. These differences may arise from differences in the presynaptic inputs carrying binocular input to PV and SST neurons and/or differences in the postsynaptic mechanisms that shape synaptic integration and input-output transformations in PV and SST neurons in binocular V1. To investigate this possibility, we next determine the intrinsic and synaptic properties of PV and SST neurons in bV1.

### Intrinsic and synaptic properties of PV and SST neurons in bV1

In V1, feedforward thalamic inputs primarily carry contralateral eye signals, whereas interhemispheric callosal projections from the opposite V1 are thought to primarily carry ipsilateral responses ^13,32,42,43^. We therefore asked whether both PV and SST interneurons in bV1 received interhemispheric input from the contralateral bV1 (Figure 4A). To investigate this we injected channelrhodopsin2 (ChR2) coupled to YFP into V1 in the right hemisphere in mice expressing tdTomato in either PV or SST interneurons (Figure 4B,C). In coronal slices containing V1, we identified bV1 in the left hemisphere by the presence of dense YFP-labelled callosal axons ^43–45^ and obtained whole-cell recordings from visually identified layer 2/3 PV and SST interneurons in bV1 based on tdTomato fluorescence (Figure 4C). PV interneurons in bV1 exhibited markedly faster membrane dynamics than SST neurons. On average, the membrane time constant of PV interneurons was approximately three-fold smaller than that of SST neurons (Figure 4D), suggesting synaptic potentials will rise and decay faster in PV compared to SST interneurons, limiting the temporal window for integration ^46^. PV interneurons also showed reduced sag in response to hyperpolarizing current steps compared to SST interneurons (Figure 4E), consistent with higher expression of the hyperpolarization-activated conductance I_h_ in SST compared to PV interneurons. In line with this, SST interneurons had more depolarised resting membrane potentials than PV neurons (PV vs SST (mean±sd): −71±6 mV vs – 66±4 mV, KS test, p=0.009). PV interneurons also exhibited significantly lower input resistance (Figure 4E, right) and required larger depolarising current injections to elicit spiking than SST interneurons (PV vs SST (mean±sd): 225 ±112.2 pA vs 90.4 ±46.2 pA, t-test, p<0.0007), consistent with greater intrinsic excitability in SST compared to PV interneurons.

**Figure 4:**
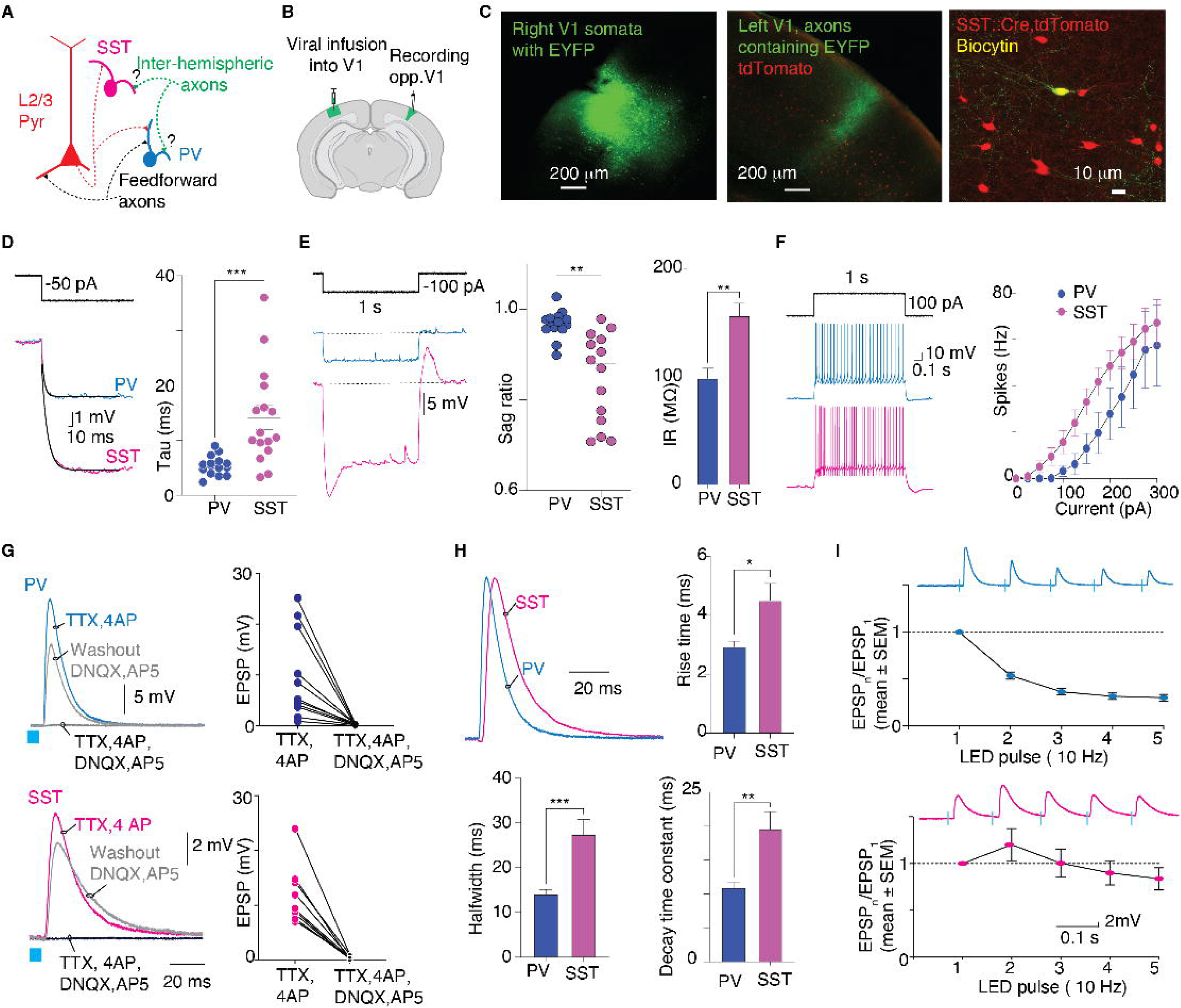
Intrinsic properties and interhemispheric inputs to PV and SST neurons in the binocular zone of V1. (A) Diagram of known synaptic feedforward and feedback connectivity in V1 (Pyr,red) to PV (blue), and SST (magenta) with putative interhemispheric connections (green). (B) Schematic of coronal section of the mouse brain showing viral infusion (AA1-ChR2) into the right V1 and recording from post-synaptic targets in left V1 in PVCre-tdTom and SSTCre-tdTom mice. (C) Left, Image of the injection site showing EYFP positive somata expressing ChR2. Middle, Image of interhemispheric axon terminals (green) in V1 and cell bodies of SST neurons (red). Right, Image of SST neurons expressing tdTomato (red) and biocytin recovery of SST neuron filled during patch clamp recording (yellow). (D) Left, Voltage response to −50 pA current injection in a PV (blue) and SST (magenta) interneuron fitted with a single exponential (black). Right, comparison of membrane time constant (tau) between PV and SST neurons (t-test). (E) Left, Voltage response to a 1 second long −100 pA current pulse in a PV (blue) and SST (magenta) interneuron. Resting membrane potential indicated by the dotted line. Middle and right: Sag ratio and input resistance (IR) in PV (blue) and SST (magenta) neurons (t-test). (F) Left, Voltage traces of PV and SST neurons spiking to long depolarizing current injections. Right, F-I curve showing spike rate (mean±sem) versus depolarizing current step amplitude in PV (blue) and SST (magenta) neurons. (G) Synaptic response generated by optical activation (470 nm, 2 ms pulse) of callosal projecting axon terminals in PV (top) and SST (bottom) in the presence of TTX and 4-AP in control and after application of AMPA (DNQX) and NMDA (AP5) receptor antagonists. (H) Scaled traces of optically evoked EPSPs in SST and PV neurons showing differences in the time-course; optically evoked EPSP rise time (top right), half width (bottom left) and decay time (bottom right) in PV and SST neurons (t-test). (I) Repeated optically evoked EPSPs (10 Hz, 5 pulses) in PV neurons showed synaptic depression (top, blue), whereas SST neurons showed moderate facilitation (bottom, magenta).

To isolate direct inputs from ChR2-expressing callosal axons, we blocked action potentials with tetrodotoxin (TTX) and enhanced neurotransmitter release by applying a low concentration of 4-aminiopyridine (100 µM) to block potassium channels. Brief 470 nm LED pulses (2 ms) evoked excitatory postsynaptic potentials (EPSPs) in all PV (n=37/37) and SST (n=24/24) neurons recorded in bV1. These responses were glutamatergic, as they were completely blocked by AMPA and NMDA receptor antagonists (Figure 4G). At matched light intensity (3mW), mean EPSP amplitudes were comparable in PV and SST neurons (PV vs SST (mean±sd): 11.6±1.92 mV vs 11.6±1.7 mV; P>0.05). Despite similar amplitudes, callosal EPSPs in PV neurons exhibited significantly faster kinetics, including shorter rise times, smaller half-widths, and faster decay times compared to SST interneurons (Figure 4H). The faster EPSP kinetics in PV compared to SST interneurons are consistent with their shorter membrane time constant (Figure 4D). In addition, a lower NMDA receptor contribution to excitatory synapses onto PV compared to SST interneurons may contribute to differences in callosal axon evoked EPSP kinetics ^47^. Similar to other work on the temporal dynamics of excitatory synaptic input to PV and SST interneurons^48–51^, callosal input to PV neurons showed synaptic depression during repetitive stimulation (10 Hz, 5 pulses), whereas callosal inputs to SST interneurons showed moderate facilitation (Figure 4I). Together, these data show that both PV and SST interneurons receive direct glutamatergic callosal input from the opposite V1, but have distinct intrinsic and synaptic properties and input-output transformations.

### Biophysically constrained modelling of temporal dynamics in PV and SST interneurons

To test how these cell-type-specific differences in intrinsic and synaptic properties in PV and SST neurons influence temporal dynamics, we simulated PV and SST activity using morphologically realistic, biophysically constrained models (Methods). To evaluate how local excitatory input shapes PV and SST activity, we drove excitatory synapses in each model with a time-varying presynaptic firing-rate profile matched to that observed experimentally in pyramidal neurons during binocular stimulation. Thus, PV and SST models received identical, temporally matched in vivo-like excitatory input. Importantly, the excitatory synaptic dynamics and synaptic distributions to the PV and SST models were constrained to those previously reported for excitatory connectivity in V1 ^47,50^, consistent with the difference in synaptic dynamics to PV and SST neurons observed during callosal input (Figure 4I).

During matched excitatory drive both PV and SST interneuron models closely tracked the time course of excitation (Figure 5A,B). Firing in the PV model peaked earlier than in the SST model (PV model 132 ms vs SST model 162 ms), consistent with the faster membrane time constant and synaptic kinetics in PV compared to SST neurons (Figure 4). This observation contrasts with our *in vivo* measurements, where on average, SST units peaked before PV units. In addition, no obvious post-peak decline in SST responses was observed in these simulations in contrast to our observations *in vivo* (Figure 2). These simulations therefore suggest that differences in intrinsic properties and synaptic dynamics on their own are insufficient to account for the divergent temporal responses of PV and SST neurons observed *in vivo* during binocular visual input.

**Figure 5.**
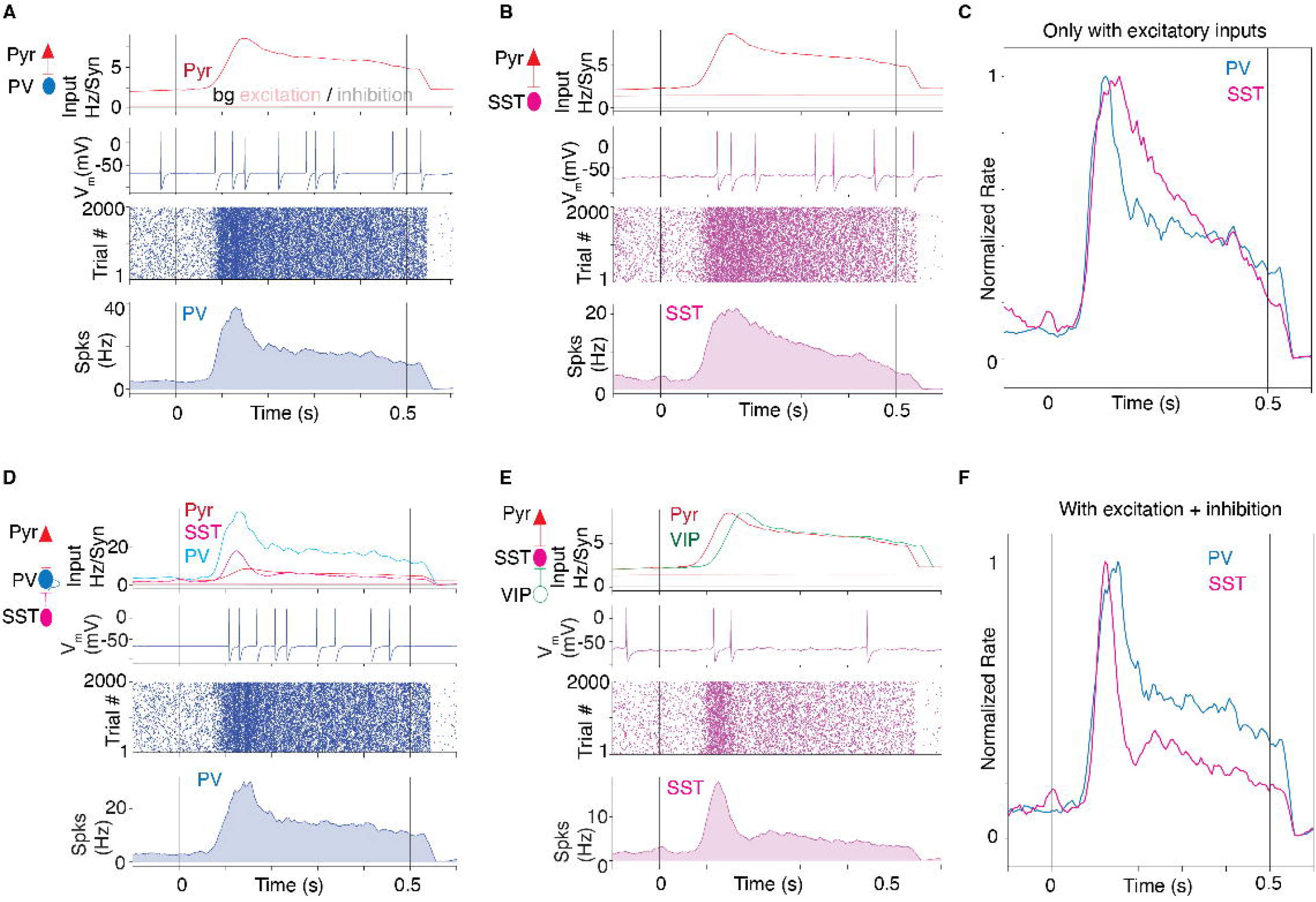
Response dynamics of biophysical models of PV and SST to multi-synaptic excitation and cell-type specific inhibition. (A) Spike responses of a biophysical model of a PV model to temporally constrained excitatory synaptic inputs emulating experimentally measured visual response in pyramidal population. Top, Time course of excitatory input rate (red) emulating visual response of pyramidal population and background (bg) inputs. Second, spike output of a PV (blue) model on a single trial. Third, raster for all trials (n=2000). Bottom, average spike rate. (B) Same as in A for the SST model (magenta). (C) Overlaid PV (red) and SST (magenta) model responses (average spike rate) during matched excitatory drive. (D) Same as in A, with addition of inhibitory inputs to the PV model. (E) Same as in B, with addition of inhibition to SST model. (F) Overlaid PV (red) and SST (magenta) model responses following addition of inhibition to PV and SST models during matched excitatory drive.

We next tested whether introducing interneuron-class-specific inhibitory circuit motifs could reproduce the key features of PV and SST *in vivo* dynamics. As PV neurons receive inhibition from SST and PV neurons ^24,50^, we tested whether adding SST-mediated inhibition to PV interneurons improved the match to our *in vivo* observations. Adding SST-mediated inhibition to PV neurons shifted the PV response profile, reducing the peak firing rate and delaying PV peak firing (compare Figure 5A and D, peak latency with only excitation vs with additional inhibition: 132 ms vs 152 ms). These simulations suggest that SST-mediated inhibition of PV neurons can delay the peak response of PV relative to SST firing, consistent with our *in vivo* observations.

To evaluate the mechanisms that could account for the rapid, post-peak decline in SST firing observed experimentally, we added inhibitory input to the SST model, emulating the VIP to SST inhibitory motif ^50,52^. This inhibitory input truncated the rising phase of SST spiking, advanced SST peak latency (excitation alone vs additional inhibition: 162 ms vs 122 ms) and produced a pronounced post-peak suppression (Figure 5E). The resulting SST response profile was qualitatively similar to that observed *in vivo* (Figure 5F, compared with Figure 2A bottom), with SST responses peaking before PV responses. Together, these simulations indicate that interneuron-class-specific inhibitory circuit motifs are required to reproduce the distinct temporal dynamics of PV and SST interneurons observed *in vivo* during binocular integration.

## Discussion

Normalization is a canonical computation observed across sensory cortices and is thought to emerge from circuits operating as an inhibition-stabilized network (ISN)^6,16^. Whether PV or SST interneurons contribute to ISN-like inhibitory dynamics remains unclear ^5^. Some modelling and experimental studies indicated that PV interneurons are the primary mediators of inhibitory stabilization ^16,53,54^, whereas others implicate an important role for SST interneurons, particularly during strong sensory input ^10,38,55^. In cortical models operating as ISNs, inhibition closely tracks excitation ^11,15,17,33^ and causes sublinear response summation ^11,18^, contributing to normalization of population responses. As described here, we find that the temporal dynamics and binocular summation in PV interneurons were broadly consistent with this inhibitory profile, whereas SST interneurons exhibited distinct temporal dynamics and summation. Notably, these divergent response dynamics only emerged during strong visual drive mediated by contralateral and binocular stimulation; a regime when ISN-like inhibitory dynamics have been proposed to normalize population response^11,15,17,33^. These results indicate that PV, but not SST neurons have ISN-like properties during binocular integration.

Previous studies indicate that sublinear summation in pyramidal neurons during binocular input requires inhibition ^12–14^. As we found that both PV and SST interneurons are binocular, our data suggest that both interneuron populations are likely to contribute to sublinear summation in pyramidal neurons during binocular visual input. While an earlier two-photon calcium imaging study reported that approximately 70% of PV neurons in bV1 are binocular ^28^, we found that almost all (96%) of optically-tagged PV units are binocular. The higher proportion of binocular PV units in our study may reflect methodological differences, as calcium imaging can underestimate spiking ^56^. In contrast, we found that just over half of the SST population were binocular (56%). This observation is surprising given our *in vitro* recordings indicated that all PV and SST neurons received callosal input, which has been shown to play a critical role in generating binocular responses in pyramidal neurons ^32,43^, and could suggest the percentage of binocular SST neurons observed in our *in vivo* experiments is underestimated. This may result from weaker excitatory drive to SST compared to PV neurons. Consistent with this, PV interneurons receive visual input from multiple sources, including thalamocortical afferents, layer 4, and layer 2/3 pyramidal neurons ^57^, whereas SST neurons are driven primarily by surrounding layer 2/3 neurons ^37,38^.

During binocular stimulation, PV interneurons exhibited sublinear response summation, whereas SST neurons showed linear or supralinear summation. Consistent with these findings, a recent *in vitro* study showed that the higher NMDA receptor contribution to excitatory synapses onto SST neurons compared to PV neurons supports supralinear summation of excitatory inputs to SST neurons ^47^. The slower membrane time constant of SST neurons will also increase the time window for synaptic integration, potentially enhancing NMDA receptor activation and supralinear synaptic integration. In addition, the higher input resistance and a more depolarized resting membrane potential of SST compared to PV interneurons are likely to work together to support the more effective input-output transformation observed these cell (Figure 4F) ^58,59^. These different synaptic integration and input-output profiles are broadly consistent with our observed differences in binocular response summation in PV and SST interneurons.

Differences in intrinsic and synaptic properties, however, were insufficient to explain the observed temporal dynamics of PV and SST neurons *in vivo*. When PV and SST models received temporally matched excitatory drive, both interneuron models tracked excitation, with responses in the PV model peaking earlier than in the SST model. These finding contrasted with our *in vivo* measurements, in which SST responses peaked earlier than PV. Introducing SST to PV inhibition delayed the PV peak response, supporting the idea that SST-mediated inhibition of PV neurons can shape the time course of PV responses.

Additionally, SST units exhibited a pronounced post-peak suppression in our stimulus conditions. A previous study using full-field flashes has shown similar rapid transitions in SST responses compared to the more sustained response of PV neurons (see ^40^). This post-peak reduction in SST activity could arise from targeted inhibitory input onto SST neurons ^55,60,61^ and/or a reduced excitatory drive ^62–64^. Adding inhibitory input to SST neurons in our model, which may arise from VIP neurons^52^, decreased the time to peak of SST responses and produced a post-peak suppression qualitatively similar to that observed *in vivo*. Further experiments will be required to determine the relative contribution of inhibition versus activity-dependent reductions in excitation, for example via pre-synaptic GABA_B_ receptors ^63^, in shaping SST temporal dynamics *in vivo*.

How do these inhibitory dynamics relate to normalization during visual processing? PV interneuron dynamics closely tracked the pyramidal neuron population during incoming visual drive. This is likely the case as work in V1 indicates that both PV and pyramidal neurons receive shared feedforward, recurrent input ^37,57^. In addition, in bV1 both pyramidal neurons ^32^ and PV (this study) receive callosal excitatory inputs. These shared input sources provide a circuit basis for PV activity to covary with excitatory population activity^33^. Because PV interneurons primarily target the somato-axonal compartment of pyramidal neurons, they can effectlively regulate spike output during incoming sensory drive. Temporally coactive and scaled somatic inhibition from PV interneurons is therefore well positioned to provide an ISN-like inhibitory mechanism for normalizing population responses ^9^.

In contrast, SST interneurons exhibited distinct temporal dynamics and a summation profile that diverged from the canonical inhibitory profile predicted by ISNs. Nevertheless, surround suppression, which is thought to be mediated by SST neurons, is often viewed as a normalization-like operation ^38,40^. During surround suppression, spiking responses of pyramidal and PV neurons are reduced by spatially extended uniform stimuli. In contrast, SST neurons scale their response with the stimulus size in a way that has been proposed to explain surround suppression^37^. In our paradigm, dichoptic binocular stimulation engages interconnected V1 circuits in both hemispheres, which may be analogous, in circuit terms, to broader recurrent network engagement produced by a large uniform stimulus. In this context, enhanced binocular response summation in SST interneurons could serve to enhance surround suppression ^20,38^. Surround suppression is thought to reduce redundant amplification of uniform stimulus features in recurrent cortical circuits ^65^. Similarly, enhanced binocular responses in SST neurons may operate to reduce redundant amplification of visual signals carrying similar stimulus features from the two eyes. Consistent with this possibility, recent imaging findings indicate that congruent and incongruent binocular stimuli enhance and reduce SST responses, respectively ^66^. Future experiments will be required to determine the spiking dynamics and response summation of SST neurons during binocular conflict.

Together, our findings reveal interneuron-class-specific temporal dynamics and response summation during binocular integration. Future work will be needed to determine how PV and SST dynamics generalize across different behavioural states and whether disruption of these dynamics contributes to disorders of sensory integration, as occurs in autism ^67^.

## Methods

### Animals

All procedures were conducted in accordance with the guidelines approved by the Animal Ethics Committee of the Australian National University and Monash University. Transgenic mouse lines: PVCre-tdTomato, SSTCre-tdTomato, PVCre-ChRN2, and SSTCre-ChRN2. We used PVCre-tdTomato and SSTCre-tdTomato lines for viral injections to map interhemispheric connections.

### Surgical procedure for viral injections

We injected viral construct AAV1-hSyn-ChR2(H134R)-eYFP-WPRE.hGH (10-fold dilution; Addgene, titer ≥10^13^) into the right V1 of PVCre-tdTomato and SSTCre-tdTomato mice. Viral constructs were injected using glass pipettes (Drummond) with a tip diameter around ∼15 µm. Injection pipettes were backfilled with mineral oil and front-loaded with viral suspension and fitted to a nanoinjector (Nanoject II, Drummond). Animals were placed inside a chamber to induce anaesthesia with isoflurane (3.5% in oxygen), then mounted on a stereotaxic frame with continued anaesthesia with isoflurane (1-1.5% with oxygen) delivered through the nose cone. Throughout the surgery, animal body temperature was maintained at 37°C using a servo-controlled heating blanket (Harvard Instruments, USA). Eye gel was applied over the eyes to prevent drying. Meloxicam (0.5mg/ml, 5mg/kg) was given for pain management.

A small craniotomy, 0.5 mm diameter was made over the binocular zone of the primary visual cortex (bV1) in the right hemisphere using the following stereotaxic coordinates: 0.5 mm anterior, 3.0 mm lateral to Lambda. Four viral injections of volume 50 nL were made at depths of 100, 200, 300 and 400 µm from the cortical surface over a period of 5-10 minutes between each depth (injection rate: 37.5 nL per min). After the final injection, the pipette was left in place for an additional 10 minutes to allow the virus to diffuse before being slowly removed from the brain using the micromanipulator. Following viral injection, the scalp incision was closed, Meloxicam (0.5mg/ml, 5mg/kg) was given for pain management and mice were allowed to recover in pre-warmed cage. Upon full recovery, mice were transferred to their home cage and monitored daily. 4-5 weeks after viral injection, acute brain slices were prepared for electrophysiological recordings.

### Surgical procedure for in vivo recordings

Adult post-critical-period male and female PVCre-ChRn2 and SSTCre-ChRn2 mice (8-14 weeks) placed in a chamber briefly to induce anaesthesia using isoflurane (3.5% isoflurane with oxygen) and mounted on a stereotaxic frame (1-1.5mg/kg) and maintained under anaesthesia with intraperitoneal injection of urethane (1-1.5 g/kg body weight, 10% w/v in water) and sedative chlorprothixene (5mg/kg,10% w/v in water). Atropine (0.3 mg/kg) was given subcutaneously to reduce secretions. Throughout the surgery, animal body temperature was monitored with an anal probe and maintained at 37°C using a feedback-controlled heating blanket (Harvard Instruments, USA). The level of anaesthesia was regularly checked with hind paw withdrawal and corneal reflex. Ophthalmic lubricant gel was applied over both eyes to prevent drying. Whiskers were trimmed. Scalp hair was trimmed, then removed completely using a hair removing cream. Bupivacaine (5mg/mL) was injected under scalp to achieve local anaesthesia. Scalp skin was removed with scissor and skull was exposed. Skull surface was irrigated with normal saline and dried using absorbent tissue. A craniotomy (∼ 2 mm diameter) was made over the left primary visual cortex with the centre of the craniotomy over the bV1 (0.5 mm anterior, 3.0 mm lateral to lambda). A recording well was built around the craniotomy using dental cement and exposed brain area was covered with normal saline. A custom-made head-bar was glued to the rostral part of the skull and stabilized with dental cement.

### Visual stimulation

Sinusoidal drifting gratings were generated using the PsychToolbox presentation software for MATLAB. Visual stimuli were displayed on two 8.1” x 6.5” retina displays (resolution, 2048 x 1536) to stimulate each eye independently via a haploscope. The haploscope mirrors (silver coated, 15 cm X 6 cm) were mounted on a custom designed holder placed ∼2 mm in front of the snout. A thin black piece of carboard was placed in the middle of the haploscope, extending over the snout to ensure monocular stimulation. Black cardboard was also placed on either side of the head to blocking lateral visual fields. Full-field sinusoidal grating stimuli were displayed at 100% contrast, with a temporal frequency of 2 Hz and spatial frequency of 0.07 cycles/degree, drifting in 12 equally spaced directions (0°–330°). The visual stimulus was presented for 0.5 s (1 cycle) interleaved by randomized inter-trial intervals 1 s to 1.25 s of blank (grey) screen with the same mean luminance as the stimulus. Stimuli were randomized for direction and eyes.

### In vivo electrophysiology

Extracellular single unit activity was recorded using a Neuronexus 16 channel silicon array probe with 4-shanks (spacing between channels: 100 µm) mounted on a metal pole fitted to a micromanipulator (Sutter Instruments). The array was inserted vertically into bV1 using stereotaxic coordinates and arterial landmarks. The probe was slowly inserted to a depth 450 µm from the pial surface. Electrical signals were acquired using a 16-channel Omniplex recording system (Plexon, Dallas, TX, USA), bandpass filtered (250 to 5000 Hz) and sampled at 40 KHz. Location of electrode in bV1 was checked online from population spike response to monocular stimulation to drifting sinusoidal gratings (0:90:330) with spatial frequency of 0.04 cycles/degree.

### Optical tagging for PV and SST neurons

PV and SST neurons expressing channelrhodopsin were activated by blue light via an optic fibre (Thorlabs, 200 µm, 0.39 NA) coupled to a LED (470 nm) positioned 0.2 mm from the recording array over the craniotomy just above the brain surface. Blue light pulses (1.4 mW, 10 trials, 50 ms long pulses with 1 s intertrial interval) to activate PV and SST neurons were presented at the beginning and at the end of visual stimuli triggered using a digital output (PCIe-6321, National Instruments).

### Acute brain slice preparation

Under isoflurane anaesthesia, animals were perfused trans-cardially with ice-cold cutting artificial cerebrospinal fluid (aCSF) containing (in mM): 118 C_5_H_14_ClNO (choline chloride), 2.5 KCl, 2.5 CaCl_2_.2H_2_O, 5 MgSO_4_.7H_2_O, 1.2 NaH_2_PO_4_, 26 NaHCO_3_ and 10 glucose, bubbled with carbogen (95% O2, 5% CO2), pH=7.4, and osmolarity of 310-320 mOsm. The animal was then decapitated and the brain rapidly removed and submerged in ice-cold and carbogenated (95% O_2_ and 5% CO_2_) cutting aCSF. Acute coronal slices (350 µm) containing visual cortex were prepared using a vibratome (Leica VT1200S, Leica, Germany) and transferred to a holding chamber containing pre-warmed aCSF of the following composition (mM): 126 NaCl, 3.5 KCl, 1.2 CaCl_2_.2H_2_O, 5 MgSO_4_.7H_2_O, 1.25 NaH_2_PO_4_, 26 NaHCO_3_ and 10 glucose, pH 7.4, osmolarity 300-320 mOsm and continuously bubbled with carbogen. Slices were allowed to recover for 30 minutes at physiological temperature (36°C) and subsequently stored at room temperature (24-26°□C). After the 30-minute recovery period, slices were transferred one at a time to a submerged recording and imaging chamber perfused with carbogenated and pre-warmed aCSF containing 1mM MgCl_2_.6H_2_O instead of 5 mM MgSO_4_ perfused at ∼5 ml/min atC36°C. For whole-cell recordings, glass pipettes (Drummond) were filled with (in mM): 130 K-gluconate, 7 KCl, 10 HEPES, 4MgATP, 0.3 Na2GTP, 7 Na_2_Phosphocreatine, and 0.3% Biocytin, pH=7.3 and 280-290 mOsm. Pharmacological reagents were dissolved in water or DMSO, diluted in ACSF, and perfused into the recording chamber. The concentration of reagents for bath applications were: Tetradotoxin (1 µM; Hello Bio), 4-aminopyridine (100 µM; Sigma), 6,7-dinitroquinoxaline-2,3-dione (DNQX,20 µM;Tocris), Amino-5-phosphonovaleric acid (APV, 100 µM; Tocris), and Gabazine (5 µM, Tocris).

### In vitro electrophysiology

Slices were visualized under an upright microscope Olympus-BX50WI fitted with infrared differential interference contrast optics using a 60x water immersion objective (Olympus). LED light sources (470 nm, and 565 nm, ThorLabs) and filter cubes (ThorLabs) were used for epifluorescence detection of eYFP and tdTomato fluorescent proteins. Neurons were imaged with a video camera (SciTech). Electrical signals were acquired with a Dagan BVC-700A amplifier, digitized using an ITC-18 A/D converter (InstruTECH, Germany), and recorded and analysed using Axograph X (Axograph Scientific, USA). Electrical signals were low-pass filtered at 10 kHz and digitized at 50 kHz. Intracellular voltage was not corrected for liquid junction potentials. In each slice, the right V1 (injection site) was visualized to verify somata expressing eYFP and the left V1 (projection site) for eYFP fluorescence due to interhemispheric axon/terminals. Recordings targeted the fluorescent band in the left V1 representing callosal projections to neurons in binocular V1. In transgenic mice (PVCre-tdTomato and SSTCre-tdTomato), PV and SST neurons were targeted based on tdTomato fluorescence excited by 565 nm LED light. We recorded from neurons in layer 2/3 within 400 µm from the pia. To characterize intrinsic electrophysiological properties, neurons were injected with 500 ms long current pulses of different magnitudes (−100 pA to 400 pA, 25 pA steps). After characterising intrinsic electrical properties, we characterised excitatory post-synaptic potentials (EPSPs) generated by activation of callosal axons expressing channelrhodopsin (ChR2). ChR2 was excited using 470 nm LED light (1 ms, 3 mW) at 0.1 Hz with the recorded neuron at the centre of the circular field of illumination (∼330 µm in diameter). Following this, slices were perfused with aCSF containing tetrodotoxin (to block action potentials) and 4-AP (to enhance neurotransmitter release) to isolate monosynaptic responses. In a subset of experiments, we elicited LED trains (5 pulses at 10 Hz) to investigate short-term synaptic plasticity. To verify that responses were synaptic we bath applied glutamatergic receptor antagonists (DNQX and AP5).

### Extracellular spike waveform detection and classification

Single units were isolated offline using clustering algorithm Wave-Clus in MATLAB^68^. Spike waveforms were detected with a negative threshold four-times the estimated standard deviation of the background noise and minimum refractory period of 3 ms. Spike waveforms were subsequently analysed for trough to peak distance and spike half-width using custom written codes in MATLAB. PV or SST neurons expressing ChR2 were identified as optically tagged units if they responded to 470 nm light pulses within <10 ms, and exhibited a statistically significant increase in firing during the optical stimulation window. Significance was assessed by Receiver Operating Characteristics (ROC) analysis by comparing baseline firing rates (50 ms before light stimulation) and during the light pulse (50 ms) with shuffled data (1000 times). Units with ROC values >0.9 were classified as optically tagged. Units that did not show light evoked responses and had trough to peak distances of > 0.6 ms and spike half-widths > 0.25 ms were characterised as putative pyramidal units (see Figure 1D) ^21^.

### Detecting visually responsive units

All in vivo electrophysiological data were analysed using custom written codes in MATLAB. Visually responsive units were identified by a significant increase in the spike rate within 300 ms from stimulus onset using a baseline 500 ms prior to visual stimulation and confirmed using bootstrap resampling. For bootstrap resampling, 1000 re-samplings with replacement were made and two-tailed assessments of significance was calculated from observed rates to the 95th percentile of the extremes (highest 97.5th percentile or lowest 2.5th percentile) of the surrogate distributions. Units responding to both contralateral and ipsilateral eye stimulation were classified as binocular.

We estimated the ocular dominance index (ODI) based on contralateral and ipsilateral eye responses to drifting gratings using the following formula:

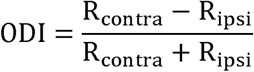

where R_contra_ and R_ipsi_ are mean spike rates for drifting grating stimulus presented to the contralateral and ipsilateral eyes, after subtracting the baseline firing rate. Contralateral and ipsilateral dominance are indicated by 1 and −1, respectively. A cell equally responsive to either eye has an ODI = 0.

The orientation selectivity index (OSI) was estimated as 1-circular variance of the peak firing rate across all orientations. An OSI of 1 indicates a case where a cell responds exclusively to one orientation, while an OSI of 0 indicates a cell responding equally to all orientations.

### Spike response to visual stimulation

For each unit, the spike counts were binned in 5 ms bins. The average spontaneous spike rate was subtracted from the instantaneous spike rate for each unit. To normalize spike rates, we estimated the baseline subtracted average spike rate for the recording duration excluding optical tagging trials. The baseline subtracted instantaneous spike rate for each unit was divided by its average spike rate to generate the normalized spike response. For each unit, the monocular sum of responses was estimated from the arithmetic sum of contralateral and ipsilateral responses. For each unit, spike responses were smoothed using a five-bin moving mean (movean, MATLAB). Response properties, including peak latency, rise time, and post-peak decline, were then measured from the smoothed response of each unit individual. Because smoothing can alter absolute latency estimates, we used the same smoothing window across all units and cell types for comparisons. Results were similar when analysis was performed without smoothing.

The summation index was calculated as:

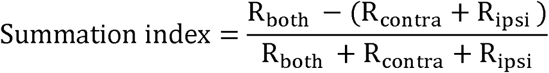

where R_both,_ R_contra_ and R_ipsi_ indicate responses to the stimulation of both eyes, contralateral and ipsilateral eye, respectively. A summation index of zero indicates linear summation, positive values indicate supra-linear, while negative values indicate sublinear response summation.

### In vitro analysis

Intracellular voltage responses were analysed using Axograph, with plots generated in Prism (GraphPad, USA). The membrane time constant was estimated from the initial voltage response to 1 second −50 pA current injections. The voltage response from 0.5 ms to 150 ms was fitted with one or more exponentials with added constant using a simplex algorithm to minimise the sum of squared errors. The membrane time constant was identified as the slowest time constant. Sag in the voltage response to long hyperpolarising current pulses was quantified as the ratio of the peak membrane potential at steady state (0.8 to 1s) divided by the peak voltage during the initial (0-0.2s) response during 1 second −100 pA current injections. Input resistance (R_in_) was calculated from the steady-state membrane potential (V_m_) response to a −100 pA current pulse (I) and calculated using Ohm’s law: R_in_ = V_m_/I. Synaptic kinetics were estimated from an average of 15 trials.

### Immunohistochemistry

During whole-cell patch recording, neurons were filled with biocytin and neuronal morphology was recovered by immunohistochemistry. After recording, slices were transferred to a 16 well plate containing 4% paraformaldehyde (PFA) in 0.1 M phosphate buffered saline (PBS; pH 7.2) and incubated for 1 h at room temperature for rapid fixation and then stored at 4° C for overnight incubation. On the following day, PFA fixed slices were resuspended in 0.1 M PBS and rinsed 3 times for 15 min. Slices were subsequently submerged in blocking buffer: 1%BSA, 0.05% saponin, and 0.05% sodium azide in 0.1M PBS, and incubated on a shaker for 1h at room temperature. To label biocytin, Alexa Fluor 647 bound to streptavidin (1:2000, Invitrogen) was added to the blocking buffer and slices were incubated overnight at room temperature. On the following day, the blocking buffer was removed, slices were washed with 0.1 M PBS (3 times for 15 min), and then mounted on a glass slide with a mounting medium (Vectasheild, Vector Laboratories, USA). Immunolabeled neurons were imaged under an upright fluorescence microscope (Zeis; Axioscope II, USA).

### Morphologically realistic SST and PV neuron models

Models of PV and SST neurons were based on models taken from the Allen Brain database (#484635029 for PV and #485466109 for SST) with passive membrane properties after incorporation of active conductances adjusted to match experimental observations (Figure 4; Table 1). The membrane capacitance was set to 1 μF/cm^2^ ^69^. Simulations were performed using NEURON ^70^ with a time step of 0.025 ms.

Active conductances in the PV model were: (i) Na_T(a)_, Na_P_, K_D_, K_v2like_, K_v3-1_, K_T_, SK, Ca_HVA_, Ca_LVA_, and I_M_ in the soma, (ii) K_T_, K_D_, K_v2like_, K_v3-1_, SK, Ca_HVA_, and Ca_LVA_ in the axon, and (iii) K_v3-1_, and I_M_ in the dendrites. Active conductances in the SST models were: (i) Na_T(a)_, Na_P_, K_D_, K_v2like_, K_T_, SK, Ca_HVA_, Ca_LVA_, I_M,_ and I_H_ in the soma, (ii) Na_T(a)_, K_T_, K_D_, K_v2like_, SK, Ca_HVA_, and Ca_LVA_ in the axon, and (iii) I_M_, and I_H_ in the dendrites. See Table 2 for details.

### Synapse locations and properties

For the PV model, both excitatory and inhibitory synapses were placed in a distance-dependent manner based on relationship: −0.015x+4.25 synapses/µm (set to 0 if negative), where x denotes the distance from soma (in µm). For SST model, excitatory and inhibitory synapses were distributed uniformly along the dendrites at a density of 2 synapses/μm ^47^. For excitatory synapses, NMDA to AMPA ratio was set to 1.5 for the SST model, whereas the NMDA conductance was omitted from PV model ^47^.

Excitatory transmission was mediated by AMPA and NMDA receptors, and inhibitory transmission by GABA_A_ receptors, as in our previous models ^71,72^. AMPA receptors were modelled as a point process using a double exponential conductance profile ^73^, in Eqns. 1, 2,

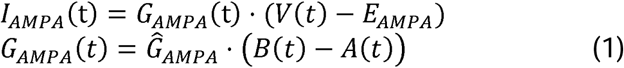

where *I_AMPA_* is the current produced by the synaptic population of AMPA receptors; *E_AMPA_* is the reversal potential of the receptor; *V* is the membrane potential; *G_AMPA_* is the conductance of the receptor population, with peak *Ĝ_AMPA_*.

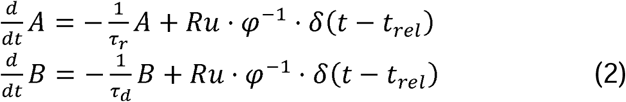

where 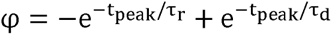 and *t_peak_* = *τ_r_τ_d_*/((*τ_d_* – *τ_r_*)log(*τ_d_*/*τ_r_*)). Variable *A* models the rising component of the conductance, with time constant *τ_r_*; variable *B* models the decaying component of the conductance, with time constant *τ_d_*; *t_rel_* is the time of a release event; *R* and *u* are the synaptic resources and efficacy, respectively, at each *t_rel_*, which evolve according to short-term plasticity as described below; *t_peak_* is the time to peak of the conductance and *φ* is a normalization factor such that *G_AMPA_*(*t_peak_* + *t_rel_*) = *Ĝ_AMPA_* assuming only one release event and *Ru* = 1. The subscript AMPA for *A*, *B*, *φ*, *t_peak_*, *τ_r_*, *τ_d_*, *R* and *u* was dropped to improve readability. These variables and parameters are independent from those for the NMDA receptor, described next.

The NMDA receptor was modelled in a similar manner, but incorporated the voltage dependence due to the magnesium block ^74^, as shown in Eqn. 3 and replacing the AMPA subscript replaced by NMDA,

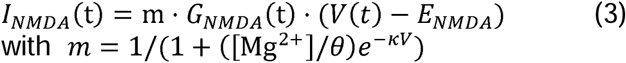

where *I_NMDA_* is the current; *E_NMDA_* is the reversal potential of the receptor; *V* is the membrane potential; m is the magnesium block gating variable; *θ* is the scaling factor for the extracellular magnesium concentration [Mg^2+^]; *κ* is the slope of the voltage dependence of magnesium block.

For all synapses mediated by AMPA/NMDA receptors, we used the same reversal potential *E_AMPA_* = *E_NMDA_* =0 mV, the same variables *R* and *u* for short-term plasticity, and a fixed ratio for synaptic conductance *Ĝ_NMDA_* = 1.5 *Ĝ_AMPA_*. For the AMPA receptor, the transient dynamics were governed by rise time constant *τ_r_* = 0.2 ms, decay time constant *τ_r_* = 1.7 ms for SST and *τ_r_* = 0.2 ms, *τ_d_* = 1.7 ms for both PV and SST models, and the following for the NMDA receptor ^75^: *τ_r_* = 0.3 ms, *τ_d_* = 43 ms for both cells. The other NMDA parameters were: [Mg^2+^] =1 mM, *θ* = 3.57 mM, *κ* = 0.062 mV^-1^.

The GABA_A_ receptor dynamics were modelled using Equations 1 and 2 replacing the AMPA subscript by GABA_A_. The reversal potential was 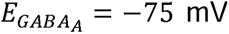, and the time constants were *τ_r_* = 0.5 ms, *τ_r_* = 5.5 ms for connections to both SST and PV cell types.

### Synaptic short-term plasticity

Model AMPA/NMDA and GABA_A_ synapses incorporated short-term presynaptic plasticity (STP). Specifically, excitatory synapses from PN to PV were depressive and those from PN to SST were facilitatory. None of the synapses had long-term plasticity. STP was incorporated into all excitatory synapses using the Tsodyks–Markram formulation ^76,77^, wherein repeated presynaptic spikes modulated the maximal synaptic conductance by a fraction of the synaptic efficacy (*U*) and synaptic resources available for transmission (R):

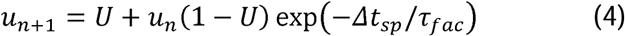

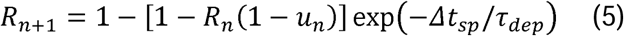

where τ_fac_ and τ_dep_ are the time constants of depression and facilitation respectively, and *U* can be interpreted as the initial release probability. *R* and *U* are updated at the time of each presynaptic spike, and *Δt_sp_* is the interval between the current and the previous spike. The tuned short-term plasticity parameters were as follows: (U, *τ_dep_*, *τ_fac_*) = (0.4,0,70) for PN-SST, and (0.8,100,0) for PN-PV synapses.

### Modeling afferent inputs

PV and SST models had two types of synapses: background synapses and task synapses. Accordingly, two types of afferent input trains fed those synapses, ‘background’ inputs representing the various connections that were provided to all cells via background synapses, and ‘task’ inputs representing the visual response modelled from *in vivo* measurements.

### Background inputs

Background synapses received independent Poisson spike train inputs that were tuned to reproduce a pre-task baseline membrane potential ∼– 60 mV ^47^. For the PV model, background inhibitory and excitatory synaptic inputs were 0.75 and 0.025 Hz, respectively. For SST model, inhibitory and excitatory inputs were 1.5 Hz and 0.05 Hz, respectively.

### Task (Visual) inputs

Task-related inputs representing visual responses were modelled as additional Poisson spike trains delivered to task synapses and superimposed on background activity. For each interneuron type, the number of task synapses was derived from Allen Institute connectivity data: Depending on the simulation (see Results), PV cells received on average 713 excitatory afferents from pyramidal neurons and in some simulations additional 52 and 70 inhibitory inputs representing SST and PV inputs, respectively. SST cells received 435 excitatory afferents from PNs and in some simulations additional 95 inhibitory synaptic inputs representing VIP inputs.

PN inputs to task synapses of PV and SST models were generated by providing the experimentally measured PN-firing rate curve (Figure. 2) as input to an inhomogeneous spike-train generator^47^. Independent spike trains generated from this process provided the excitatory task inputs to each model. Thus, PV and SST models received temporally matched excitatory input derived from the experimental PN firing rate profile. In simulations incorporating inhibitory circuit motifs, the PV model additionally received inhibitory inputs from SST and PV populations, whereas the SST model received inhibitory inputs from VIP population. VIP input firing rates were assumed to track the PN drive, with a 30 ms delay^78^. Mean PV and SST model responses were obtained by averaging across 2000 simulation trials.

Once background inputs had been tuned, task inputs were added. Conductances of individual model synapses for both types of inputs were sampled from lognormal distributions with the means reported below and a standard deviation equal to 0.33 times the mean ^79^. Mean synaptic weights and background firing frequencies were tuned to reproduce the baseline and peak firing rates observed experimentally. For the PV model, the mean AMPA and GABA conductance were 0.5 nS and 0.4 nS, respectively. For the SST model, the mean AMPA and GABA conductances were 1.5 and 3.0 nS, respectively.

### Quantification and statistical analysis

Statistical analyses were performed in MATLAB and GraphPad. Sample sizes, including animal counts, cell/unit counts, and the statistical tests used are reported in the Methods, Results and figure legends. Statistical significance was classified using the asterisk (*) system based on P values as follows: ns, not significant, P > 0.05, * P <= 0.05, ** P <= 0.01, *** P <=0.001, **** P <= 0.0001.

## Author Contributions

M.B.P, and G.J.S conceptualized the project, designed the experiments, and interpreted the data. M.B.P performed the in vivo and ex vivo experiments and data analysis. H.B. and D.D. generated the models performed the simulations under the supervision of S.S.N. S.G. and E.A. contributed code for in vivo data acquisition, curation and analysis. M.B.P drafted the manuscript and all authors edited and approved the final version of the manuscript.

## Data availability

The data supporting the findings of this study are available from the corresponding authors upon reasonable request.

## Code availability

Custom-written MATLAB scripts used for data acquisition and analysis are available from the corresponding authors upon reasonable request.

