## Supplemental Table 1 and 2 for "Cell-type specific inhibitory dynamics shape binocular response normalization in visual cortex"

**Table 1. Passive properties of the PV and SST models**

|  | **PV Model** | **SST Model** |
| --- | --- | --- |
| RMP (mV) | -71 | -65 |
| Tau (ms) | 5.6 | 15.8 |
| Sag ratio | 0.99 | 0.85 |
| Rin (MOhm) | 95 | 152 |

**Table 2. Passive and active parameters of the SST and PV cell***

| PV neuron | | | |
| --- | --- | --- | --- |
| Property | Soma | Axon | Dendrite |
| *Passive Parameters* | | | |
| Gpas (S/cm²) | 17.5e-5 | 17.5e-5 | 17.5e-5 |
| Epas (mV) | -71 | -71 | -71 |
| Cm (µF/cm²) | 1.0 | 1.0 | 1.0 |
| Ra (Ω·cm) | 33.5 | 33.5 | 33.5 |
| *Active Parameters; Channel densities* (S/cm²) | | | |
| I_h_ | - | - | - |
| Na_p_ | 2e-3 | - | - |
| Na_Ta_ | 0.2 | - | - |
| K_d_ | 6e-2 | 4.12e-3 | - |
| Kv_2_like | 1e-2 | 5.26e-2 | - |
| Kv_3_ | 9e-2 | 0.765 | 0.71 |
| K_T_ | 1e-3 | 2.05e-4 | - |
| I_m_ | 1e-3 | - | 2e-3 |
| SK | 1e-5 | 1.0e-5 | - |
| Ca_HVA | 3.6e-3 | 9.55e-5 | - |
| Ca_LVA | 5e-3 | 3.8e-3 | - |
| SST neuron | | | |
| Property | Soma | Axon | Dendrite |
| *Passive Parameters* | | | |
| Gpas (S/cm²) | 5.8e-5 | 5.8e-5 | 5.8e-5 |
| Epas (mV) | -65 | -65 | -65 |
| Cm (µF/cm²) | 1.0 | 1.0 | 1.0 |
| Ra (Ω·cm) | 150 | 150 | 150 |
| *Active Parameters; Channel densities* (S/cm²) | | | |
| I_h_ | 2e-3 | - | 2e-3 |
| Na_p_ | 1.3e-3 | - | - |
| Na_Ta_ | 0.1 | 0.1 | - |
| K_d_ | 1.8e-2 | 5.6e-3 | - |
| Kv_2_like | 2e-4 | 0.1 | - |
| Kv_3_ | - | - |  |
| K_T_ | 1e-4 | 0.37 | - |
| I_m_ | 5e-5 | - | 1e-2 |
| SK | 3e-5 | 1.62e-6 | - |
| Ca_HVA | 2.5e-3 | 3e-5 | - |
| Ca_LVA | 1e-3 | 3.5e-3 | - |

* Ion channel mechanisms and initial parameters obtained from Allen Cell types Database. Channel mechanisms were retained and selected parameters and conductances were adjusted to match experimental measurements.
